# A simple *ex vivo* bladder infection model permits study of host-pathogen interactions in UTI

**DOI:** 10.1101/2024.12.23.630047

**Authors:** Rixa-Mareike Köhn, Méline Durand, Laura Ramirez Finn, Ariana Costas, Bill Söderström, Livia Lacerda Mariano, Molly A Ingersoll

## Abstract

Urinary tract infections (UTI) are one of the most common infections, worldwide. To understand mechanisms of UTI pathogenesis and find new treatments, researchers often use animal models, such as mice or rats. However, studying certain phenotypes in animals can be difficult. Additionally, using animals in research comes with significant administrative and ethical challenges. To address these challenges, we developed a simple, reproducible, and cost-effective model to study UTI using donated mouse bladder tissue that would otherwise be discarded. This model allows researchers of all experience levels to study interactions between the host and pathogen in a controlled environment. We tested uropathogenic *E. coli* colonization and invasion of isolated urothelial sheets from 30 minutes to 24 hours, finding that bacterial burden in our *ex vivo* model was comparable to *in vivo* UTI mouse models. To optimize reproducibility, we tested multiple variables, including technical parameters, such as incubator conditions, and biological factors, such as biological sex or prior pregnancy in the donor mouse. This method offers several advantages, including assessment of early host-pathogen interactions, immune cell uptake of bacteria, the impact of age and sex of donor animals in infection, and diverse bacterial strains, mutants, or treatments. In addition, in some countries, sharing material recovered from animals sacrificed for other reasons does not require additional ethical approval by the receiving laboratory, providing a resource for labs without access to animals and reducing administrative burden. Given the breadth of the model with respect to sex, age, mouse and bacterial strain, and the ability to test any parameter that can be included in a 96-well plate, we believe this model will be useful to UTI researchers, with potential application beyond infection or even beyond the bladder to other tissues.

## Introduction

To investigate pathogen interactions and host immune responses, investigators often rely upon animal models. Studying host-pathogen interactions *in vivo* provides the opportunity to observe and manipulate local and systemic responses to infection. There are limitations to *in vivo* studies, however, including the assessment of very early or rapid events. In addition, the use of animal studies necessitates consideration and application of the three Rs of animal experimentation: to Replace animal models, Reduce the number of animals used, and/or Refine models to minimize unnecessary or unjustified stress and suffering (Lewis, 2019). While application of the 3Rs to animal research should be a part of designing an experimental study, replacing *in vivo* models can be challenging. One approach to reduce animal numbers, while maintaining the complexity of an organ, is to use organs recuperated from animals used for other purposes in an *ex vivo* model. *Ex vivo* models have the advantage of providing multiple samples from the same animal to test different conditions and reduce biological variability. Examples include *ex vivo* skin explant models to test topical products or lung slice models to investigate mycobacterial infection (Eberlin et al., 2020; Molina-Torres et al., 2020). Explant models also exist to observe host-pathogen interactions at mucosal surfaces, such as the bladder urothelium (Wieser et al., 2011). In one such model, an explanted mouse bladder is sandwiched in a custom organ chamber, which is the same size as a standard microscope slide, to facilitate live imaging. The apparatus is sealed and can be connected to syringe pumps to add or collect fluids or materials during experimentation. This model was used to investigate very early pathogen interactions with host urothelium and local immune responses. A second *ex vivo* model, in which a 2 mm^2^ piece of bladder is slowly stretched by microtranslators in a custom aluminum incubation chamber over a microscope slide, permits live imaging and was used to capture uropathogenic *E. coli* (UPEC) kinetics during infection (Justice et al., 2004). Both models are well-adapted to microscopy applications, however, they both require a custom set-up that may not be widely available to investigators, limiting their application. Thus, a more universally accessible *ex vivo* bladder model would support investigation of the dynamics of urinary tract infection (UTI) and the host response, while reducing the number of mice used for experimentation.

The bladder urothelium, which lines the urethra and bladder, constantly interacts with commensal and pathogenic microbes residing in or entering the urinary tract (Khandelwal et al., 2009). The most prevalent pathogen, UPEC, causes 70-80% of community-acquired UTI and a majority of hospital-associated infections (Flores-Mireles et al., 2015). UTI affect more than 400 million people, worldwide, every year and, recurrent UTI arises in approximately half of those who have experienced a first infection (Yang et al., 2022). Alarmingly, the number of multi-drug resistant uropathogens is increasing rapidly, underlining the need for new, non-antibiotic-based approaches to treat acute and recurrent UTI (Totsika et al., 2011). To develop new approaches, it is necessary to identify bacterial targets involved in pathogenesis or host targets mediating an efficacious immune response. *Ex vivo* infection models may provide new ways to test therapeutic strategies, while reducing reliance on animals.

Building on a protocol to separate the urothelium from the bladder muscle wall (Lu et al., 2019), we established an *ex vivo* infection model in peeled urothelial sheets to test bacterial invasion, colonization, and local host response under different conditions without the need for custom equipment. We established this model in line with the 3R principles by using bladders donated by our colleagues from naïve mice sacrificed for other reasons. We optimized this protocol to use a single bladder for up to four different experimental conditions. We found that *ex vivo* infection with the commonly used UPEC strain UTI89 and a pandemic ST131 strain, JJ2528, was comparable to published *in vivo* infection (Mora-Bau et al., 2015; Price et al., 2013). We used this model to test different parameters, including colonization in nulliparous or puerperal female mice and in male mice. We found that resident immune cells were present in the dissected tissue and took up bacteria similarly to our *in vivo* model (Mora-Bau et al., 2015; Zychlinsky Scharff et al., 2017; Zychlinsky Scharff et al., 2019). This approach was easily mastered by new investigators, including those with limited laboratory experience, demonstrating the ease and reproducibility of the model. In sum, we developed a cost-efficient method to investigate early host-pathogen interactions during UPEC infection, which could be adapted to other organs, such as the kidney or prostate, or other mucosal surfaces.

## Results

### Ex vivo urothelial sheets are colonized similarly to bladders infected in vivo

We first wanted to determine whether an explant model would reliably reflect bacterial colonization observed in a widely used female mouse UTI model (Hung et al., 2009; Zychlinsky Scharff et al., 2017). Using an institute-wide email list, we requested that any laboratories sacrificing naïve mice of any sex or strain to contact us if they would be willing to share tissue. We established a consistent source of bladders, dissected from naïve animals sacrificed for other reasons, which would otherwise be discarded.

Bladders from 6-14 week old naïve female C57BL/6 mice were collected within 2 hours of sacrifice, and cut longitudinally into four approximately equally sized pieces, using a scalpel and a dissecting microscope. Following a published protocol to separate bladder layers (Lu et al., 2019), we separated the luminal-facing urothelial and submucosa layers from the underlying muscle layer by gentle peeling with forceps. To maintain simplicity of the model, we did not further separate the submucosa from the urothelium and used the ‘urothelial sheets’ with the associated submucosa. Each urothelial sheet was placed in a well in row A of a sterile 96-well round bottom tissue culture plate prefilled with sterile PBS (**Figure 1A**). Once all bladder quarters were peeled and placed in the 96-well plate, urothelial sheets were infected by moving the tissue with forceps to row B of the plate containing 2-2.5×10^6^ colony forming units (CFU) of the UPEC strain UTI89-RFP-kan^R^ (Mora-Bau et al., 2015). We chose 2.5×10^6^ CFU to infect urothelial sheets as this is one quarter of the inoculum we use for *in vivo* infection, *i*.*e*., 1×10^7^ CFU (Zychlinsky Scharff et al., 2017). 96-well plates were incubated in a 37°C bacterial incubator for 24 hours. After incubation, the urothelial sheets were moved successively to rows C, D, and E, containing sterile PBS, for 5 minutes washing by gentle agitation. Finally, the quarters were homogenized in 1 mL sterile PBS, serially diluted, and plated on agar plates for quantification.

**Figure 1.**
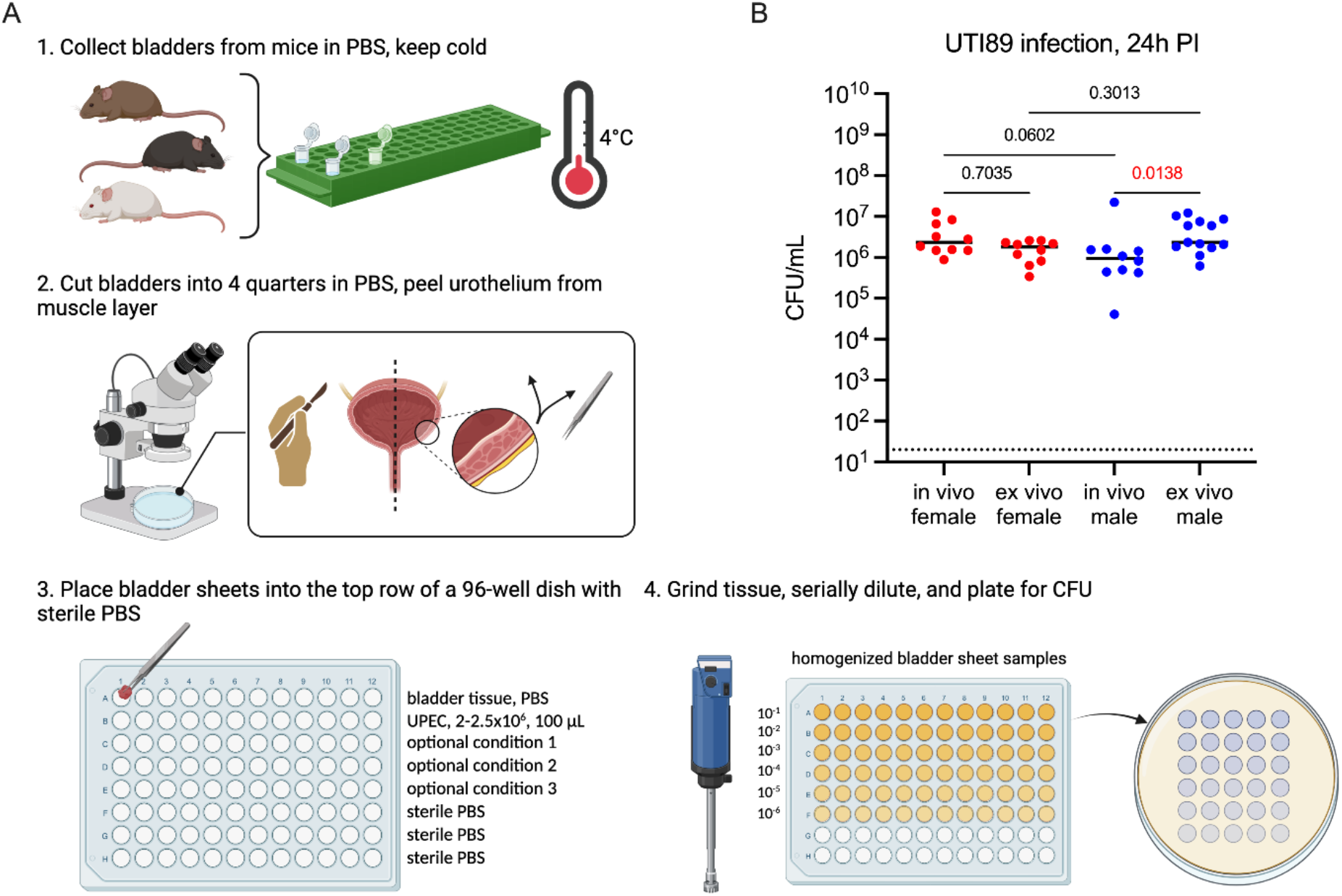
A simple *ex vivo* protocol recapitulates *in vivo* UPEC colonization. (**A**) Graphic depicts the optimized protocol to infect urothelial sheets from mice. (**B**) Graph shows colony forming units (CFU) per mL following homogenization of bladders infected *in vivo* with 1×10^7^ CFU of UPEC strain UTI89-RFP-kan^R^ or bladder sheets infected ex vivo for 24 hours with 2.5×10^6^ CFU of UTI89-RFP-kan^R^. Whole bladders or urothelial sheets were homogenized in 1 mL PBS, CFU reported as CFU/mL (equivalent to CFU/bladder in vivo, or CFU per urothelial sheet). Red denotes female mice, blue is male mice, each dot is one mouse or one urothelial sheet. Dotted line shows the limit of detection of the assay, 20 CFU. *p*-values were calculated by Kruskal-Wallis test with correction for multiple comparisons using the Dunn’s test. All comparisons made are shown and those that are *p<0*.*05* are shown in red. Experiment was performed 2-3 times with n=3-5 per experiment. (**A**) created in BioRender, https://BioRender.com/c26×567.

The median CFU (1.8×10^6^ CFU per urothelial sheet) was not statistically significantly different from the median CFU (2.3×10^6^ CFU per bladder) from *in vivo* infected female mouse bladders (**Figure 1B**). We previously reported that the immune response to UTI and outcome following infection differs starkly between female and male mice, however this is not due to differences in colonization at 24 hours post-infection (PI) (**Figure 1B** and (Zychlinsky Scharff et al., 2019)). To test whether differences in colonization exist between the sexes in our *ex vivo* urothelial sheet model, we infected male C57BL/6 mice and urothelial sheets from male C57BL/6 mice. We observed that CFU from male urothelial sheets were not different than CFU from female urothelial sheets (**Figure 1B**). We did observe that CFU from urothelial sheets were slightly higher than CFU from *in vivo*-infected male mice, however CFU in the *ex vivo* infection model also appeared to be less variable than typically observed in male *in vivo* infections (**Figure 1B** and (Zychlinsky Scharff et al., 2019)).

### Urothelial sheet infection is robust in variable conditions

To determine whether other UPEC strains behaved similarly in the urothelial sheet model, we infected urothelial sheets from male C57BL/6 mice with 2.5×10^6^ CFU of UTI89-RFP-kan^R^ or the clinically derived, multi-drug resistant, extraintestinal pathogenic *E. coli* (ExPEC) strain JJ2528 (Price et al., 2013). This strain belongs to the sequence type ST131 and is a global pandemic UTI strain with antimicrobial resistance. We observed that colonization was similar between the two UPEC strains 24 hours PI (**Figure 2A**).

**Figure 2.**
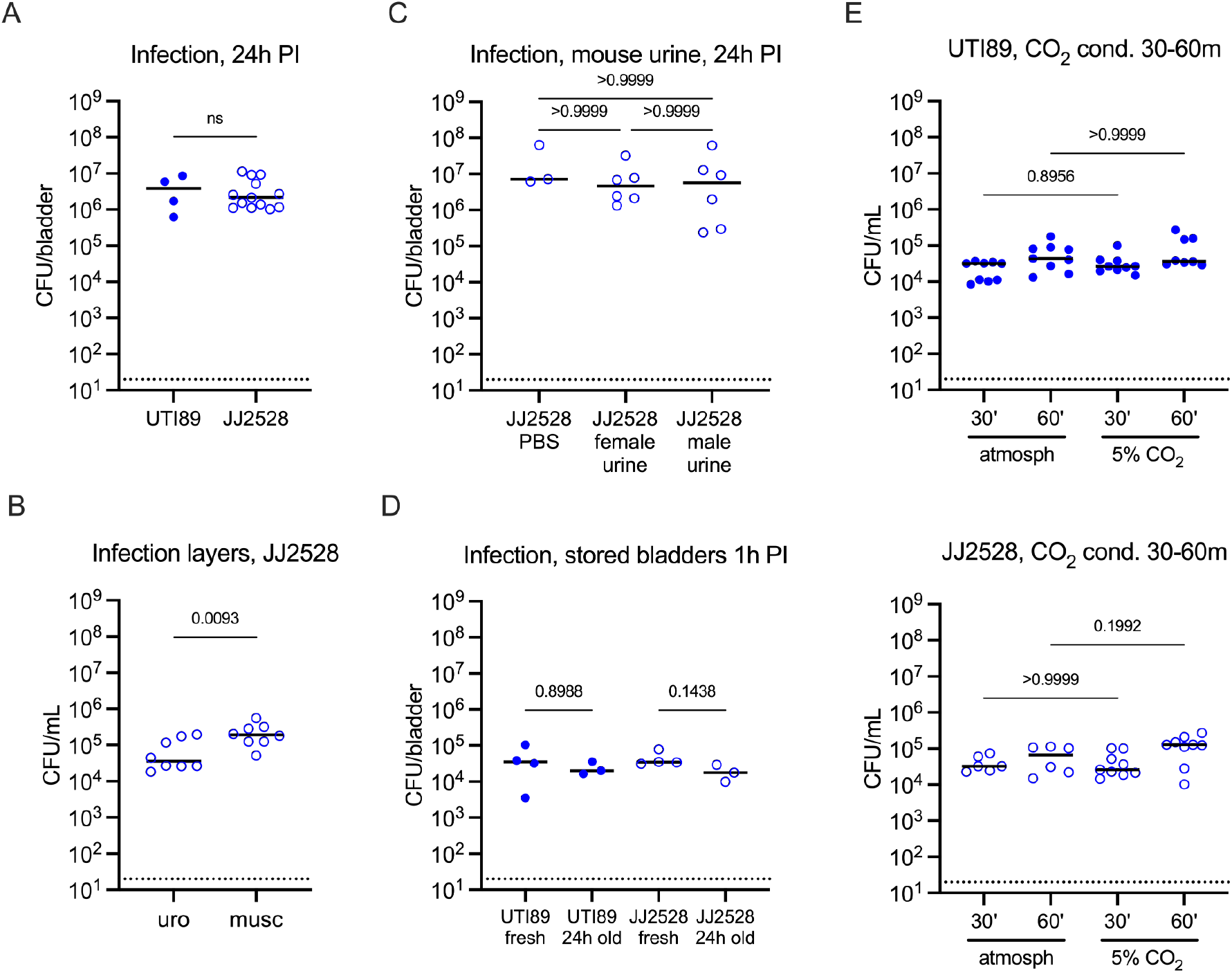
Urothelial sheet infection is robust under different conditions. Male urothelial sheets were infected with 2.5×10^6^ CFU of UTI89-RFP-kan^R^ or the pandemic ST131 strain JJ2528, as indicated. Graphs show colony forming units (CFU) per mL following homogenization, serial dilution, and plating of samples comparing the following conditions: (**A**) UTI89 versus JJ2528 at 24 hours PI, (**B**) urothelial sheets versus muscle sheets at 1 hour PI, (**C**) PBS versus mouse urine at 24 hours PI, (**D**) freshly collected versus 24 hour refrigerated bladders at 1 hour PI, (**E**) incubation in atmospheric conditions versus 5% CO_2_ at 30 and 60 minutes PI. Only male urothelial sheets were used; each dot is one urothelial sheet. Dotted line shows the limit of detection of the assay, 20 CFU. *p*-values were calculated by Mann-Whitney (2 comparisons) or Kruskal-Wallis test with correction for multiple comparisons using the Dunn’s test. All comparisons made are shown and those that are *p<0*.*05* are shown in red. Experiments were performed (**A, E**) 2-3 times, n=2-4 per experiment, (**B**-**D**) 2 times, n=2-4 per experiment.

We questioned whether it was necessary to peel the urothelium from the muscle layer to assess urothelial colonization. Therefore, we separately infected urothelial sheets and muscle from male C57BL/6 mice infected with JJ2528 and compared CFU at 60 minutes PI. Surprisingly, the muscle layer had a significantly higher bacterial burden then the urothelial sheets (**Figure 2B**). As invasion of the muscle was a result of incubating bacteria with the tissue, and is unlikely to occur *in vivo* early in infection, we continued to use peeled urothelial sheets to investigate early host urothelial-pathogen interactions.

We next considered that one major difference between *in vivo* and *ex vivo* infection was the presence of urine. Thus, we tested whether performing the *ex vivo* infection in the presence of urine would change bacterial colonization. We infected male C57BL/6 urothelial sheets with 2.5×10^6^ CFU of UTI89-RFP-kan^R^ resuspended in PBS or in fresh female or male mouse urine. We determined CFU at 24 hours PI, finding that the presence of urine did not change CFU associated with the urothelial sheets compared to infection performed in PBS (**Figure 2C**). These results suggest that the early stages of infection may not be dependent on the local nutrient microenvironment.

Given that bladder availability could not always be foreseen, we tested whether storing intact bladders impacted bacterial colonization. We used male C57BL/6 bladders donated either 24 hours prior to infection and stored at 4°C in PBS or received on the day of infection to test this condition. All bladders were peeled just before infection with UTI89-RFP-kan^R^ or JJ2528, as above. As our main objective in developing this model was to assess short term infection, and to limit the impact of further aging of the tissue outside the animal, we quantified CFU after 1 hour of incubation. We observed that preserving intact bladder tissue at 4°C for 24 hours did not impact CFU of either strain 1 hour PI (**Figure 2D**).

While optimizing our infection, we considered that our *ex vivo* model lies between *in vivo* models and cell culture models. Given that infections in *in vitro* cultured cells are typically performed in 5% CO_2_ tissue culture incubators, we tested whether infecting urothelial sheets under different incubation conditions impacted colonization. We infected urothelial sheets from male C57BL/6 mice with UTI89-RFP-kan^R^ or JJ2528 and incubated the 96-well plates in either a bacterial incubator with atmospheric CO_2_ levels or a 5% CO_2_ tissue culture incubator. We limited these infections to 30 and 60 minutes to assess short term infection in these conditions. We observed no differences between the two environments for the early time points tested (**Figure 2E)**. All subsequent experiments were incubated in a bacterial incubator with atmospheric CO_2_ levels.

### Biological sex does not impact infection of urothelial sheets

Our initial experiments showed that urothelial sheets had similar bacterial burden between female and male mice at 24 hours PI (**Figure 1**), which is in line with our published *in vivo* results (Zychlinsky Scharff et al., 2017; Zychlinsky Scharff et al., 2019). To have insight into early events during infection between the sexes, we infected urothelial sheets from female and male C57BL/6 bladders, incubating the samples for 30 or 60 minutes, before washing, homogenizing, and serially plating. In addition to UTI89-RFP-kan^R,^ we tested infection with JJ2528. Interestingly, we observed a significant increase in CFU/bladder sheet from 30 to 60 minutes of infection with both UPEC strains in female mice and only with UTI89 in male mice (**Figure 3A-B**). These results suggest that these infection conditions generally support bacterial growth, although this may not be universally true.

**Figure 3.**
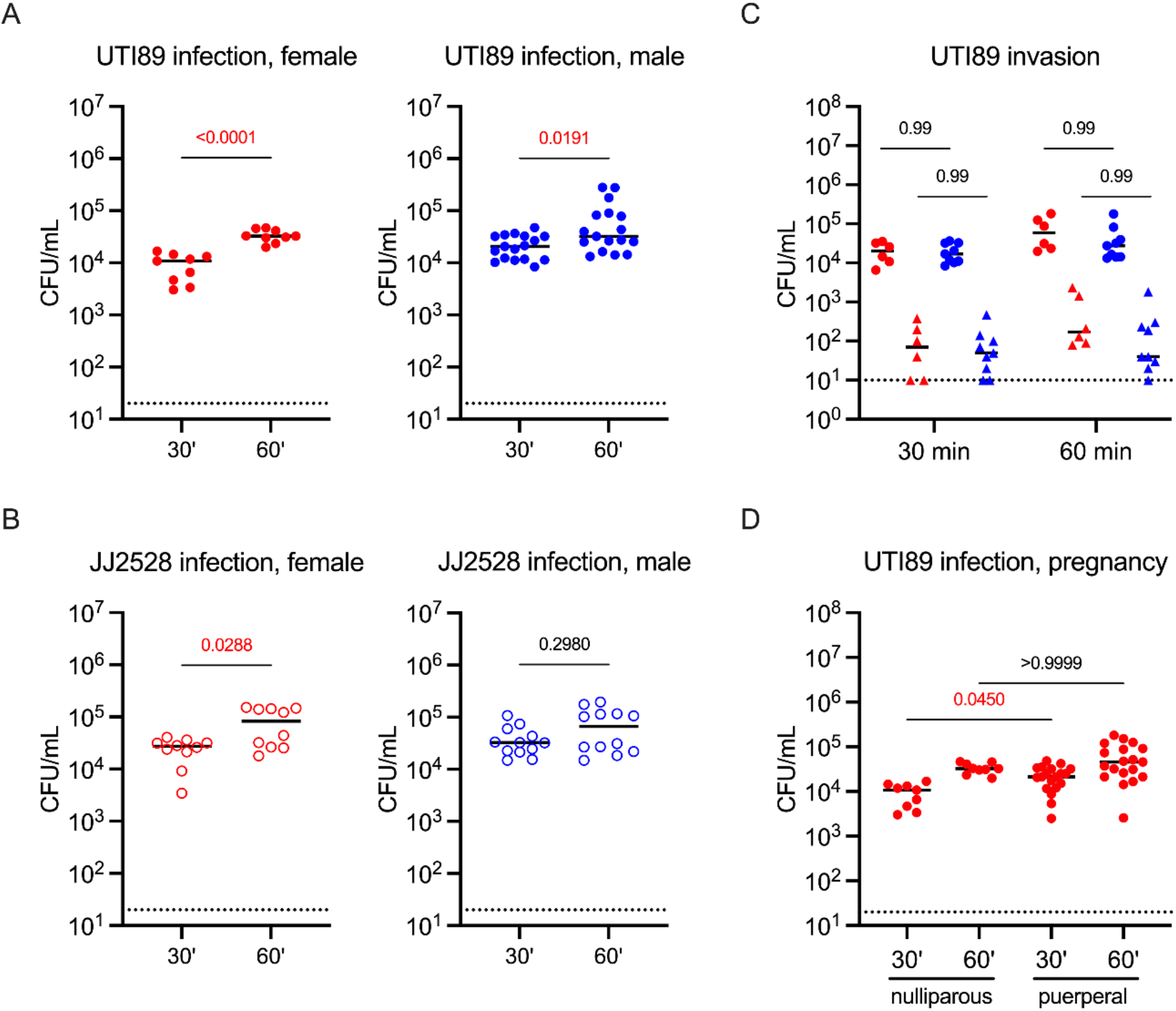
UPEC invade urothelial sheets rapidly. (**A-B**) Graphs show CFU from female and male C57BL/6 urothelial sheets infected for 30 or 60 minutes with (**A**) 2.5×10^6^ CFU of UTI89-RFP-kan^R^ or (**B**) 2.5×10^6^ CFU of UPEC strain JJ2528. (**C**) Graph shows CFU from female (red) or male (blue) urothelial sheets infected with 2.5×10^6^ CFU of UTI89-RFP-kan^R^ for 30 or 60 minutes and either mock treated (circles) or treated (triangles) with gentamycin for 60 minutes before tissue processing. (**D**) Graph shows CFU in urothelial sheets from nulliparous or puerperal mice infected with 2.5×10^6^ CFU of UTI89-RFP-kan^R^ for 30 or 60 minutes. Dotted line shows the limit of detection of the assay, 20 or 10 CFU, each dot is one urothelial sheet. *p*-values were calculated using a (**A, B**) Mann-Whitney test or (**C, D**) Kruskal-Wallis test with correction for multiple comparisons using the Dunn’s test. All comparisons made are shown and those that are *p<0*.*05* are shown in red. Data in (**A, B**) are pooled from 3-4 experiments with n=3-4 per experiment. Data in (**C**) are pooled from 2 experiments (female) or 3 experiments (male) with n=3 per experiment. Data in (**D**) are pooled from 2-5 experiments with n=1-3 per experiment.

We next considered that even though bacterial infection appeared to be similar between the sexes at these early timepoints, urothelial invasion may be different. To determine the number of bacteria that invaded, as compared to those that were attached to the surface or loosely associated with the tissue, we performed the same experiment as in Figure 3A and incorporated an antibiotic incubation step. Urothelial sheets were infected with UTI89-RFP-kan^R^. At 30 or 60 minutes PI, the sheets were moved to wells containing 100 μg/mL gentamycin (row C), or PBS as a control, for 60 minutes. Gentamycin kills extracellular bacteria, while intracellular bacteria are protected from killing at this concentration and duration of treatment. After gentamycin treatment, urothelial sheets were washed 3 times and CFU was quantified. We observed that, in line with published *in vivo* results (Schwartz et al., 2011), less than 1% of bacteria invaded the urothelial sheets (0.006% female 30 min, 0.009% female 60 min, 0.005% male 30 min, 0.006% male 60 min). Supporting that biological sex does not impact infection or invasion in this model, CFU in the control (total bacteria associated with the tissue) or antibiotic-treated (intracellular) conditions were not different between the sexes (**Figure 3C**).

From one of our collaborators, we had access to bladders from nulliparous (never pregnant) and puerperal (at term or just delivered) mice. Approximately 10% of pregnant women experience UTI, which can cause serious complications, including low birth weight, preterm labor, hypertension, and systemic infections (Szweda and Jozwik, 2016). Thus, we tested whether bacterial colonization was different between mice that had or had not experienced recent pregnancy. Using the same experimental approach, we infected urothelial sheets from nulliparous mice or puerperal mice that had delivered pups within 24 hours of bladder dissection for 30 or 60 minutes. Interestingly, bacterial colonization was significantly higher in urothelial sheets from puerperal mice at 30 minutes, but colonization differences were no longer evident at 60 minutes PI (**Figure 3D**).

### UPEC encounter specific immune cell populations in urothelial sheets

One objective in establishing this model was to observe early events in the urothelium during infection, including the response of resident immune cells. For example, we previously reported that one subset of resident macrophages, MacL, is enriched in the lamina propria (Lacerda Mariano et al., 2020). To identify the immune cells present, and determine whether specific populations were enriched in the urothelial sheets, we analyzed whole bladders, urothelial sheets, and muscle layers from female and male mice by flow cytometry. To compare cell numbers in naive urothelial sheets and muscle layers to naïve whole bladders, we peeled bladder tissues, and then processed all four pieces of urothelial sheet or muscle together, to have the equivalent amount of tissue. We observed that the number of total immune cells was similar between the urothelial sheets and the muscle layer, with each layer harboring approximately 50% of the total immune cells found in the bladder (**Figure 4A**). The total number of resident dendritic cells, which includes the cDC1 and cDC2 subsets, was similar between the urothelial sheet and the muscle layer. Interestingly, the number of cDC2 were similar between the two subcompartments, however, the cDC1 population, generally thought to favor cross-presentation of antigen to CD8^+^ T cells, was enriched in urothelial sheets compared to the muscle layer (**Figure 4B**). The number of total resident macrophages was evenly distributed between the urothelium and the muscle layer (**Figure 4C**). However, as we previously reported, MacM macrophages were 4 times more abundant in the muscle layer and MacL macrophages were enriched more than 10-fold in the urothelial sheets (**Figure 4C**) (Lacerda Mariano et al., 2020). Most resident macrophages in naïve bladders express the antigen-presenting molecule MHC II, however a small proportion of MCH II^-^ macrophages are also present (Lacerda Mariano et al., 2020). We observed that these macrophages were more abundant in the muscle layer, potentially providing a clue to their function. Monocytes were only detected in some naïve bladders as these are circulating cells that infiltrate the bladder during *in vivo* infection (Mora-Bau et al., 2015). Their distribution was similar between the layers, whereas eosinophils were more abundant in the muscle layer (**Figure 4D**). Finally, while we did not perform exhaustive phenotyping of lymphoid cells, we observed that total CD3^+^ T cells and specifically, γδ T cells, were enriched in urothelial sheets (**Figure 4E**), suggesting a tight association with the urothelium and potential surveillance of this tissue.

**Figure 4.**
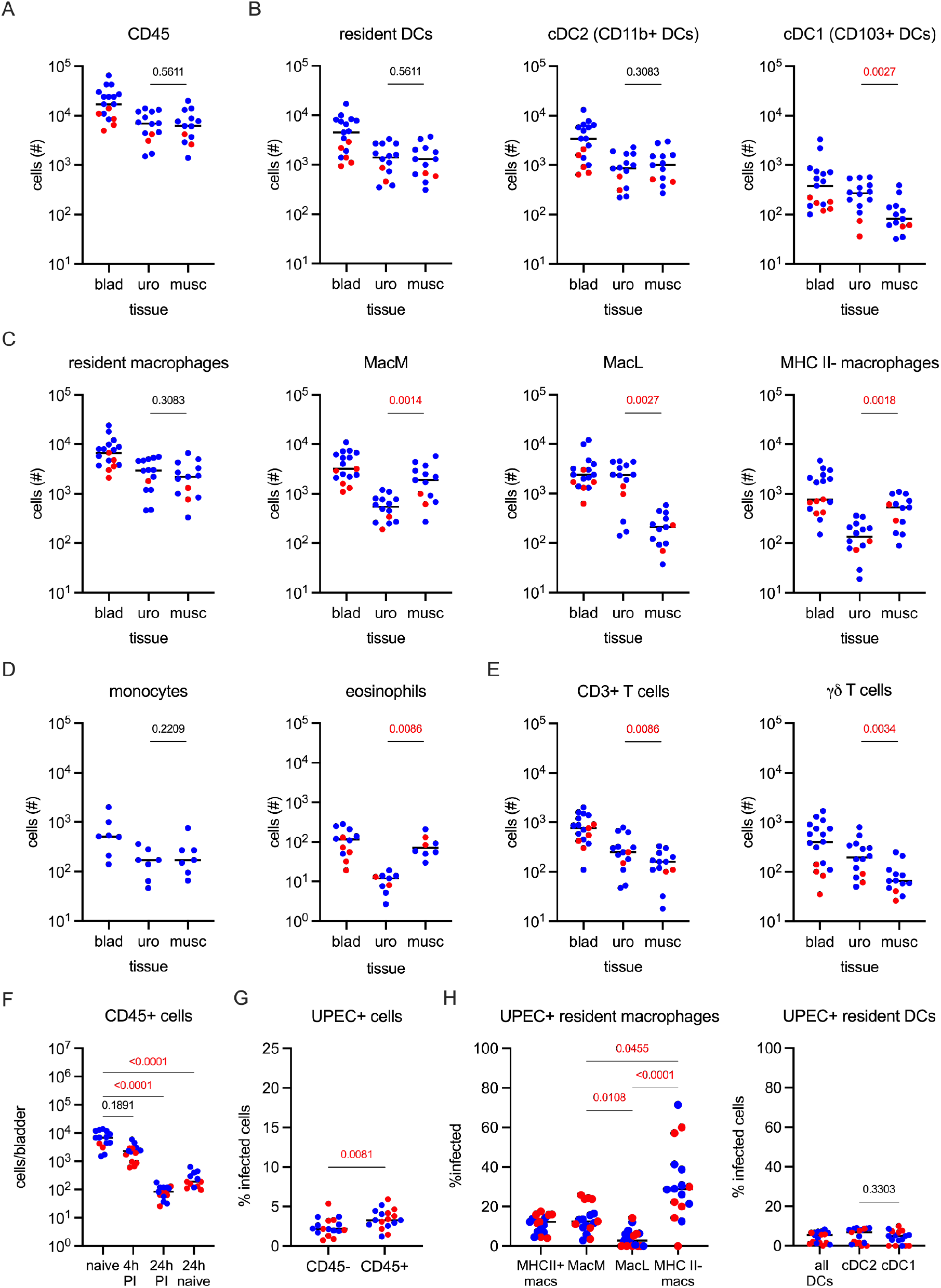
Specific immune cell populations are enriched in urothelial sheets and take up UPEC. (**A-E**) Naïve bladders, urothelial sheets, or muscle layers from naïve female and male mice were analyzed by flow cytometry. Graphs show the total number of cells of (**A**) immune cells, (**B**) resident dendritic cells and the cDC1 and cDC2 subsets, (**C**) resident macrophages and macrophage subsets, (**D**) monocytes and eosinophils, and (**E**) CD3^+^ and γδ T cells. (**F-H**) Urothelial sheets were infected with UPEC as above for 4 or 24 hours before processing for flow cytometry. (**F**) Graph shows the total number of CD45^+^ immune cells in fresh naïve bladders, bladders infected for 4 or 24 hours, or uninfected bladders incubated for 24 hours. Graphs depict the proportion of (**G**) infected CD45^-^ non-immune cells and CD45^+^ immune cells and (**H**) infected macrophages, macrophage subsets, DCs, and DC subsets. Female mice are in red, male mice are in blue, each dot is either a whole bladder, or 4 urothelial sheet quarters, or 4 muscle quarters. (**A-E, G, H** (DC graph only)) Comparisons of urothelium to muscle (paired data) were made for each cell population by Wilcoxon test and all p-values were analyzed to determine the q-values to correct for multiple testing. (**F, H** (macrophage graph only) Comparisons among infected groups were made using a Kruskal-Wallis test with a Dunn’s post-test to correct for multiple comparisons. All comparisons made are shown and those that are *q<0*.*05* are shown in red. Experiments were performed 2-6 times, n=3-9 experiments.

Having determined which immune cells were present, we next investigated bacterial colonization. We infected urothelial sheets for 4 or 24 hours observing that the total number of immune cells decreased over time (**Figure 4F**). We considered that while infection may impact the total number of immune cells, it may also be that immune cells die or leave the tissue over the 24 hour incubation period, similar to the spontaneous emigration of DCs from human explanted skin (Lukas et al., 1996). To test these possibilities, we incubated urothelial sheets for 24 hours under the same conditions as infection, finding that the total number of immune cells was significantly reduced even in the absence of bacteria (**Figure 4F**). As this observation complicated interpretation of bacterial uptake, we assessed the number of immune cells containing bacteria only at 4 hours. We found UPEC inside both CD45^+^ immune cells and non-immune cells (**Figure 4G**). UPEC could be found inside both resident macrophages and dendritic cells. Recapitulating our published *in vivo* findings, more MacM contained UPEC compared to MacL macrophages (both MHC II^+^) (Lacerda Mariano et al., 2020). Interestingly, MHC II^-^ macrophages while making up only a small proportion (**Figure 4C**), contained more bacteria than MacM or MacL macrophages (**Figure H**). Overall, macrophages contained more bacteria compared to the proportion of infected DCs, as we previously reported *in vivo* (**Figure H**) (Mora-Bau et al., 2015).

### Urothelial sheets can be used for live imaging

Having demonstrated that urothelial sheets can be used to assess infection by quantifying bacterial burden in different media and by flow cytometry, we next wanted to determine their amenability to imaging. Given that the urothelial sheets do not contain the highly scattering muscle layer, we anticipated that the optical properties would be optimized in our approach. Urothelial sheets were infected using UTI89-msfGFP-Spect^R^ (Iosifidis and Duggin, 2020), washed, stained with fluorescent dyes, spread on thin agarose pads, and imaged without fixation using either epifluorescence, quasi-Total Internal Reflection (TIRF) or confocal fluorescence microscopy on two standard commercial microscope set-ups from Nikon and Leica, respectively. Using low-magnification objectives (20x), we captured images containing several hundred host cells, easily identifiable by their nuclear stains (**Figure 5A**). We then imaged urothelial sheets at higher magnification (100x objectives) at 2 hours and 8 hours after exposure to UPEC, to capture snapshots of the infection cycle (**Figure 5B, C**).

**Figure 5.**
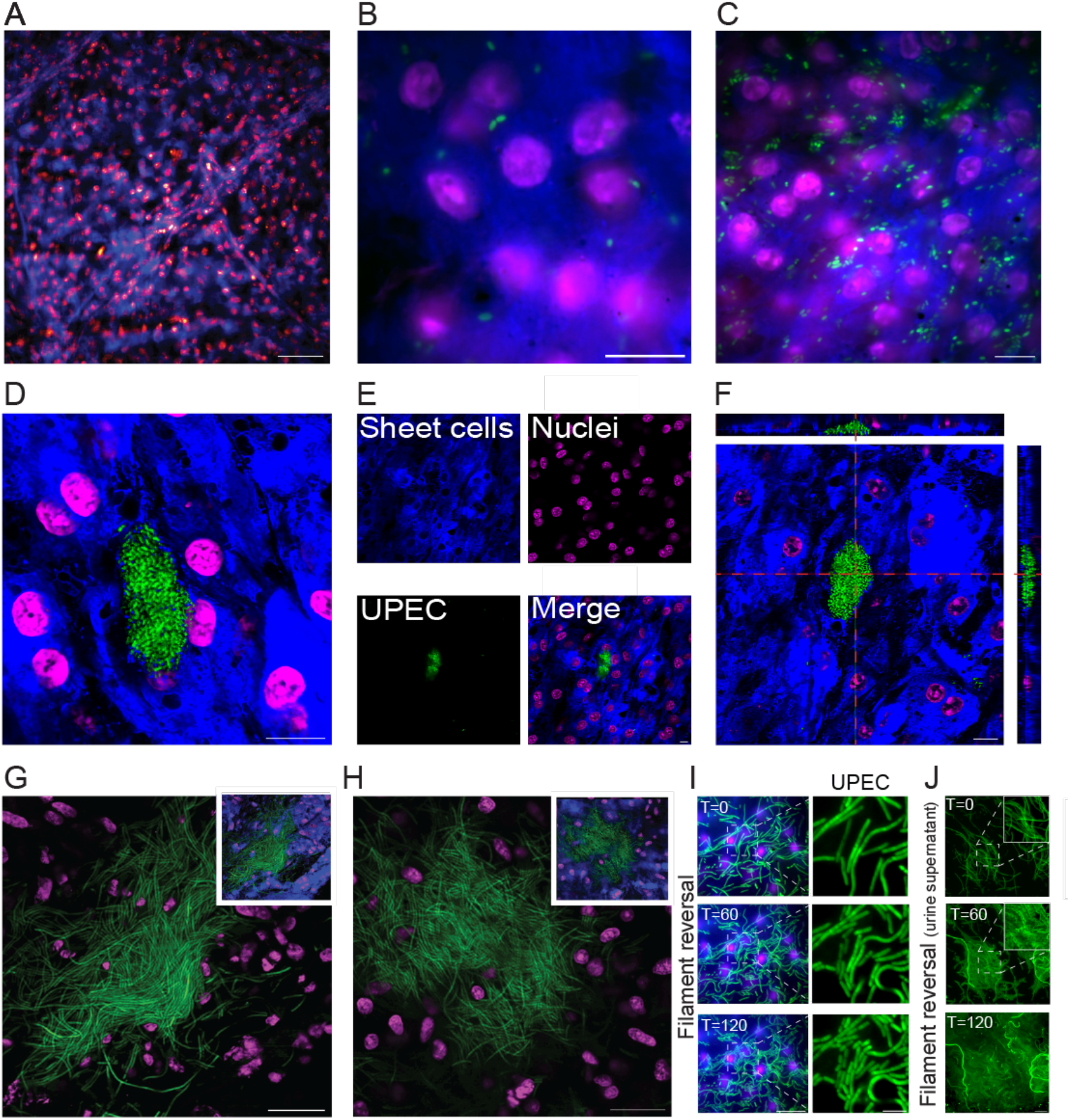
Urothelial sheets can be imaged and analyzed by live cell fluorescence microscopy. Micrographs show (**A**) low magnification (20x) epi-fluorescence image of an uninfected urothelial sheet (blue) in which more than 500 nuclei (magenta) are visible. (**B-C**) High magnification (100x) quasi-Total Internal Reflection images at (**B**) 2 hours and (**C**) 8 hours PI. (**D-F**) IBCs (green) after overnight incubation in gentamycin, in which (**F**) shows a cross section of a urothelial sheet with the IBC under the uppermost host cell layer. (**G-H**) Micrograph shows dense communities of highly filamentous bacteria after overnight incubation in human urine. Insets show images with stained urothelial cells. Micrographs show time-lapse imaging of (**I**) filament reversal on a urothelial sheet and (**J**) reversal of filaments collected from urine supernatant after overnight incubation. Time T is in minutes from first image. (**A-I**) Nuclei were stained with NucSpot650 and pseudo-colored magenta, urothelial cells were stained with CellBrite405 (blue) (**B-J**) UPEC are msfGFP (green). Scale bars: (**A**) = 50 µm, (**B-F, I** (UPEC zoom panels only), **J**) = 10 µm, (**G, H, I** (urothelial cells with bacteria panels)) = 20 µm.

We imaged infected urothelial sheets after treatment with gentamycin overnight, observing intracellular bacterial communities (IBCs)(Anderson et al., 2003), whereas only minimal numbers of extracellular bacteria were detected (**Figure 5D-E**). Confocal Z-stacks confirmed that IBCs were inside the sheets rather than above the superficial layer of bladder cells (**Figure 5F**). UPEC undergoes filamentation during UTI as a response to both the intracellular and extracellular bladder cell environment (Abell-King et al., 2022; Justice et al., 2004; Justice et al., 2006). To test whether bacteria also filament in the urothelial sheets, we treated the sheets with human urine overnight. Following overnight incubation, the sheets were washed and stained with fluorescent dyes, stretched on agarose pads, and imaged. We observed that exposure to urine induced a strong filamentation response in the bacteria, in which dense collections of highly filamentous cells, reaching lengths exceeding several hundred micrometers, could be readily resolved (**Figure 5G-H**). We assessed the viability of these filaments to determine whether they were dead or dying cells that had lost their ability to divide. Using time-lapse microscopy, we followed the reversal of filaments back to rod shape over time (**Figure 5I**). In addition, we collected and imaged the bacteria present in the urine supernatant observing large quantities of filaments, which reverted to rods within 120 minutes (**Figure 5J**), similar to published observations in *in vitro* UTI models (Andersen Thomas et al., 2012; Söderström et al., 2022). Thus, our imaging approach demonstrated that the urothelial sheet model recapitulates the main steps of the UPEC infection cycle seen in *in vivo* and *in vitro* models and supports dynamic live imaging for at least 24 hours.

## Discussion

Our objectives of this study were to develop a simple approach requiring minimal specialized equipment to investigate rapid early steps in infection, while reducing the number of mice we use for our studies. We achieved consistent results in *ex vivo* conditions, which were comparable to *in vivo* infections. The urothelial sheet was useful to assess bacterial burden, bacterial invasion, and bacterial localization. We found that generally, there was no sex difference in the early stages of infection and that recent pregnancy may increase bacterial burden, although this needs to be explored further. We also observed that different immune cell populations maintained phenotypes we previously observed *in vivo*, such as differential uptake of bacteria between macrophage subsets, and a preferential uptake of bacteria in macrophages over DCs. As immune cell numbers diminished over time, this model may be more appropriate to study immune cell responses in infections that are less than 24 hours long.

A broad range of values and measurement techniques are reported for oxygen and carbon dioxide levels in healthy bladder, and it is unlikely that they are recapitulated in the urothelial sheet model described here. We observed that different carbon dioxide concentrations did not affect bacterial burden after 30 or 60 minutes of infection. Given the flexibility of this model, urothelial sheets can be placed in any type of vessel or container for experimentation. Thus, it would be of interest to test whether urothelial sheet infection changes under diverse oxygen and carbon dioxide conditions that better reflect *in vivo* bladder physiology.

Many experiments in this study were performed in PBS, which contains no nutrients for bacterial growth. Nevertheless, we observed an increase in CFU over the initial 2.5×10^6^ inoculum in 24 hour experiments. While considered nutrient poor, UPEC can live and grow in urine (Mann et al., 2017). Our data suggest that UPEC may be able to derive nutrients directly from the urothelium, as no urine remained in the bladders prior to experimentation. Alternatively, UPEC may scavenge nutrients from dying bacteria in the well. Further experimentation is needed to address this question.

Our model has certain limitations. As this is an *ex vivo* model, there is no peripheral immune cell infiltration. This can also be considered an advantage in that it permits the isolated study of resident immune cell responses. As we observed however, specific resident immune cell populations are enriched, whereas others are present at low numbers in the urothelial sheets. Thus, the role of certain immune cells, such as the MacM macrophages predominantly found in the muscle, cannot be easily studied in this model. Another limitation is the loss of specific mechanical cues, tissue tension, and neural connections when the bladder is removed and urothelium is separated from muscle, which may impact how bacteria interact with the tissue. Despite these limitations, this model provides a robust means to investigate host-pathogen interactions. We found that the time needed to execute this model autonomously was very rapid and achievable by scientists with all levels of experience and training, including high school interns, and newly starting masters and PhD students. Additionally, we propose that this model will help reduce animal testing by using otherwise discarded bladders, which could potentially give access to animal tissues to laboratories without their own animal ethical protocol. This last point must be confirmed by researchers with their local authorities, however, in many countries, the sharing of tissues after an animal has been sacrificed is permitted. Thus, we established a simple, reproducible model to study infection in the bladder that also reduces the number of animals used for biomedical research.

## Methods

### Mice

*In vivo* animal experiments were approved by the Comité d’Éthique en matière d’Expérimentation Animale Paris Descartes (CEEA 34) under the protocol number APAFiS #34290. Mice for *in vivo* experiments in Figures 1 and 4 were obtained from Charles River Laboratories, France, and housed in the animal facilities of Institut Cochin. For this study, 6-7-week-old female and male C57BL/6J mice were used. Mouse bladders for *ex vivo* experiments were gifted to us by other laboratories of the Institut Cochin and University of Technology Sydney. All donated mice were 6-14 week old C57BL/6J from Charles River Laboratories, France or Australian Bio Resources, Australia.

### In vivo UTI model

Mice were anesthetized by intraperitoneal injection of 100 mg/kg ketamine and 5 mg/kg xylazine, catheterized transurethrally, and infected with 50 µL PBS containing 1×10^7^ colony forming units (CFU) of UPEC strains UTI89-RFP-kan^R^ or JJ2528 as previously described (Zychlinsky Scharff et al., 2017; Zychlinsky Scharff et al., 2019). Mice were sacrificed after inhaled anesthesia by cervical dislocation and infected bladders were processed for CFU determination or flow cytometry, as described below.

### Ex vivo UPEC infection

Bladders were kept in cold PBS until dissection. Bladders were dissected into equal quarters. The luminal-facing urothelial layer was separated from the underlying muscle layer using forceps under a dissecting microscope. Urothelial sheets were placed individually in the first row (A) of a sterile U-bottom 96-well plate in PBS until all samples were prepared. For infection, 2.5×10^6^ CFU of UPEC in 100 µL 1X PBS was added to row B of the plate and urothelial sheets were moved by forceps into this row. Plates were incubated at 37°C in an incubator with atmospheric CO_2_ levels unless indicated otherwise. At indicated timepoints, urothelial sheets were washed three times in sterile PBS by moving the sheets from row B to C, D, and E, every 5 minutes with gentle agitation before processing for CFU determination or flow cytometry. To quantify invaded bacteria, urothelial sheets were treated with gentamicin (100 µg/mL) after infection and before washing in a new row (*e*.*g*., row C, with washing in D, E, F), at the indicated timepoints, and incubated for an additional 60 minutes at 37°C before processing for CFU determination. In the graphs, individual dots denote a sample, which is either a urothelial quarter or four quarters to reconstitute the whole urothelium, as specified in the legends.

### Determination of bacterial burden

CFU was determined by homogenization of intact bladders (*in vivo* infections) or urothelial sheets (*ex vivo* infections) in 1 mL sterile PBS using a Precellys tissue homogenizer, serial dilution, and plating on LB agar plates with kanamycin (UTI89-RFP-kan^R^) or no antibiotics (JJ2528). The limit of detection (LOD) for this method is 20/bladder or urothelial sheet as indicated by a dotted line in graphs. For gentamicin protection assays, in addition to serial dilution, 100 µL of sample homogenate was spread on half a plate, making the LOD 10 CFU/bladder or urothelial sheet.

### Flow cytometry

Bladders, urothelial sheets, and muscle layers were processed for flow cytometry as previously described (Zychlinsky Scharff et al., 2017). Briefly, single-cell suspensions were prepared by mincing bladder pieces, followed by incubation in 0.34 Units/mL Liberase (Roche) diluted in PBS at 37°C with gentle manual agitation every 15 minutes. Intact bladders were incubated for 60 minutes, muscle layers were incubated for 30 mins, and urothelial sheets for 15 minutes. Digestion was stopped by adding FACS buffer (PBS, 2% fetal calf serum, 0.2 µM EDTA). The suspension was passed through a 100-µm filter (Miltenyi Biotec), washed, blocked with Fc block (BD Biosciences), and stained using the antibodies in **Table 1**. All samples were acquired on an BD LRSFortessa using DIVA software (BD Biosciences) and analyzed using FlowJo software (Treestar).

**Table 1:** Antibodies used in this study.

| Antibodies (anti-mouse) | Clone | Source |
| --- | --- | --- |
| CD45 | 30-F11 | BD Biosciences |
| CD11b | M1/70 | BD Biosciences |
| MHC II (I-A/I-E) | M5/144.15.2 | Invitrogen |
| CD11c | HL3 | BD Biosciences |
| SiglecF | E50-2440 | BD Biosciences |
| F4/80 | CI:A3-1 | BD Biosciences |
| CD64 | X54-5/7.1 | BD Biosciences |
| Gr1 (Ly6G/Ly6C) | RB6-8C5 | BD Biosciences |
| NK1.1 | PK136 | BD Biosciences |
| CX3CR1 | 5A011F11 | Biolegend |
| CD326 | G8.8 | BD Biosciences |
| E-cadherin (CD324) | DECMA-1 | BD Biosciences |
| CD103 | M290 | BD Biosciences |
| γδ TCR | GL3 | BD Biosciences |

### Preparation of human urine samples for filament generation

First void urine from two different donors (one female and one male) was collected in the morning and stored for at least 2 days at 4°C. Samples were centrifuged at 4500 × *g* for 8 min and the supernatant was filtered through a 0.2 μm membrane filter. The specific gravity was determined by comparison to pure water using a gravity meter. Only urine samples in the pH range 5.3-5.8 and urine specific gravity range 1.024–1.030 g ml^−1^ were used (Söderström et al., 2022). Filter sterilized samples were stored at -20°C until required. This study has human research ethics approval from the University of Technology Sydney Human Research Ethics Committee (HRCH REF No. 2014000452). All urine donors gave approval to participate in this study by informed consent under this study’s ethics approval number.

### Urothelial sheet imaging

After initial infection with UPEC strain UTI89 expressing msfGFP [pGI5, Spect^R^ (Iosifidis and Duggin, 2020)] and incubation, urothelial sheets were stained with fluorescent dyes NucSpot650 and CellBrite405 (final concentration 1:100 in PBS as per manufacturers’ recommendations, Biotium, US) for 40 minutes. Following staining, sheets were washed twice in PBS and stretched on 2% (W/V) agarose PBS pads, covered with a pre-cleaned cover glass and directly used for imaging.

Live tissue confocal imaging was performed on a Leica Stellaris confocal microscope equipped with a 63X 1.40 NA oil objective enclosed in an environmental chamber operated at 37°C and 5% CO_2_ (Oko-Lab). The fluorophores were excited by a white laser at optimized wavelengths (405, 488, and 647), and emission was collected in pre-set system optimized detector intervals for CF405, EGFP, and AlexaFluor647 to minimize channel crosstalk. Z-stacks were always acquired to validate that the IBCs were inside bladder cells. Image size was either 2048 × 2048 or 4096 × 4096, with pixel size 90 or 45 nm, respectively. The pinhole was set to 1 AU and Z-stacks were collected with software optimized step length of either 250 or 500 nm (20–60 images per stack). 3D reconstruction and deconvolution of Z-stacks were performed in the Leica LAS software and further visualized in FIJI (ImageJ). Live tissue epi-fluorescence and quasi-TIRF (Total Internal Reflection) imaging was done on a Nikon Ti2-E N-STORM microscope (with NIS v.5.30) microscope, with a 100X 1.49 NA oil objective, to increase the signal-to-noise ratio. Quasi-TIRF imaging was performed using HILO illumination, a few degrees above the critical angle, to increase the excitation volume. To support live cell imaging, the microscope was equipped with a stage top environmental chamber regulating temperature (37°C) and CO_2_ levels (5% CO_2_). The 405, 488, and 647 nm laser lines were used to excite fluorescent probes as appropriate. Acquisition times were 70-150 ms and image size 2048 × 2048 (pixel size 65 nm). Fluorescence emission was collected via emission filter sets for DAPI, FITC and ‘Normal STORM(647)’. Images were captured using a sCMOS Flash 4.0 v3 (Hamamatsu) camera. For dynamic filament reversal time-lapse imaging, one image was acquired every 10 minutes for 3 hours.

### Statistics

Data were analyzed using GraphPad Prism 10 and multiple comparisons were corrected to avoid Type 1 error. Statistical significance was determined by a non-parametric Mann-Whitney test (two group comparison, non-paired samples), a non-parametric Kruskal-Wallis test (more than two group comparison, non-paired samples) with a Dunn’s post-test, or a nonparametric Wilcoxon matched-pairs signed rank test (paired samples) and the Prism function ‘analyze a stack of p-values’. All comparisons made are shown and all corrected *p*-values (*q*-values) are indicated, *p* <0.05 was considered significant and are shown in red in the figures.

## Acknowledgements

We are exceptionally grateful to Zeyni Mansuroglu, from the lab of Florence Niedergang, Alexis Fouque from the lab of Raphael Scharfmann at Institut Cochin, and Kristine McGrath, as well as Fiona Ryan at the Ernst animal facility at University of Technology Sydney for their generosity in supplying us with bladders. We acknowledge A/Prof. Louise Cole and the Microbial Imaging Facility at University of Technology Sydney for the use of the Stellaris confocal microscope.

## Author contributions

RMK, LLM, and MAI conceived the study. RMK, MD, LRF, AC, BS, and MAI performed experiments and data analysis. RMK and MAI wrote the manuscript. RMK, MD, LRF, AC, BS, LLM, and MAI reviewed and edited the manuscript. MAI and BS obtained funding and supervised the study. Author order of shared second authorship was determined alphabetically. All authors reviewed and approved the manuscript prior to submission.

